# Ecological divergence of DNA methylation patterns at distinct spatial scales

**DOI:** 10.1101/832816

**Authors:** H. De Kort, B. Panis, D. Deforce, F. Van Nieuwerburgh, O. Honnay

## Abstract

Adaptive trait divergence between populations is regulated by genetic and non-genetic processes. Compared to genetic change, epigenetic change is unstable and short-lived, questioning its contribution to long-term adaptive potential. However, epigenetic change can accumulate over time, and may result in beneficial epigenetic memories where environments are heterogeneous. Diverging epigenetic memories have been observed across large spatial scales, and can persist through multiple generations even in the absence of the causative environmental stressor. It is unknown, however, how and to what extent epigenetic memories contribute to fine-scale population structure and evolution. Here, we performed whole genome bisulfite sequencing on 30 *Fragaria vesca* F1 plants originating from distinct ecological settings and grown in a controlled environment. Specifically, we compared methylation patterns between a steep, altitudinal gradient (<2 km) and a wide spatial gradient (>500 km). If epigenetic variation is random, arising from errors during replication and without evolutionary implications, one would expect similar amounts of epigenetic variation across populations and no spatial scale-effect. Here, we find that epigenetic memories arise even at fine spatial scale, and that both parallel and non-parallel biological processes underpin epigenetic divergence at distinct spatial scales. For example, demethylation of transposable elements consistently occurred at the large but not the small spatial scale, while methylation differentiation for most biological processes were shared between spatial scales. Acute drought stress did not result in significant epigenetic differentiation, indicating that repeated historical stress levels associated with heterogeneous environmental conditions are required for acquiring a stable epigenetic memory and for coping with future environmental change.

## INTRODUCTION

Adaptive genetic variation underlying fitness traits is considered the dominant resource upon which plants depend for evolving under environmental change. Driven by drift, mutation and migration, such genetic variation supplies populations with trait values that support local fitness and adaptive potential. However, although it is widely assumed that adaptive phenotypic variation is mainly regulated by the underlying genetic architecture, only small proportions of the total phenotypic variation observed in many species have been associated with genetic variants (Krishna Kumar *et al*. 2016; Wellenreuther & Hansson 2016; Resende *et al*. 2017). The remaining phenotypic variation, typically referred to as missing heritability, can be roughly attributed to (i) the detection limits of rare genetic variants and genetic interactions, and (ii) heritable epigenetic variation (Brachi *et al*. 2011; Miska & Ferguson-Smith 2016; Whipple & Holeski 2016; Gienapp *et al*. 2017; Banta & Richards 2018).

The role of epigenetic variation in governing adaptive evolution remains controversial, yet a growing body of literature demonstrates the ubiquity of transgenerational epigenetic transmission, and consequently considers it as a key evolutionary force (Gugger *et al*. 2016; Miska & Ferguson-Smith 2016; Lind & Spagopoulou 2018; Schmid *et al*. 2018; Zhang *et al*. 2018; Danchin *et al*. 2019). Non-random epigenetic variation has been shown to be widespread in natural populations, and to co-vary with a range of environmental stressors, including herbivory, drought, salt and temperature (Foust *et al*. 2016; Jeremias *et al*. 2018; Alonso *et al*. 2019; Gáspár *et al*. 2019). While most stress-induced methylation changes are reset to basal levels after stress relief, part of these modifications can be stably inherited across mitotic and even meiotic cell divisions (Chinnusamy & Zhu 2009; Crisp *et al*. 2016). Such a somatic or transgenerational epigenetic stress memory allows plants to cope more effectively with subsequent stresses, thus evoking considerable fitness benefits in heterogeneous environments (Crisp *et al*. 2016; Hilker *et al*. 2016). Unraveling the relative extent of intra-generational epigenetic change resulting from acute environmental stress vs. transgenerational epigenetic accumulation may contribute to our understanding of how plants rely on their epigenetic machinery for coping with environmental change.

How selection pressures affect genome-wide DNA methylation levels in natural population remains poorly explored in non-model organisms, but considerable advances have been made in *Arabidopsis thaliana*. A study involving genome-wide DNA methylation analysis of 122 *A. thaliana* accessions sampled across Eurasia showed that climate characteristics most abundantly co-varied with methylation levels of cytosines in CHH context (where H represents a G, T or A nucleotide), with CHH methylation typically indicating the involvement of transposable elements (TEs) (Keller *et al*. 2016). These findings could be related to natural selection at the level of TE-specific methyltransferase genes that facilitate demethylation of transposons when temperatures reach extreme levels, or where populations are genetically impoverished. Stress-induced demethylation of transposons boosts transposon activity and subsequent genetic change, paving the way for rapid genetic replenishment and adaptation to environmental stressors (Mirouze & Paszkowski 2011; Ito *et al*. 2016; Rey *et al*. 2016; Schrader & Schmitz 2019). The strongest associations between climate and *A. thaliana* methylation levels were, however, found in CG contexts within or near genes related to abiotic stress responses, development and reproduction (Keller *et al*. 2016). Because (i) DNA methylation has been shown to be meiotically most stable in the CG context, and (ii) the majority of reported heritable epi-mutations occurs at CG sites (Mathieu *et al*. 2007; Jiang *et al*. 2014; Stassen *et al*. 2018), climate-CG methylation associations most likely represent solid adaptive signals. A recent study corroborated the evolutionary relevance of CG methylation using a multi-generational *A. thaliana* selection experiment, demonstrating that (i) methylation of differentially methylated cytosines (DMCs) was significantly higher in CG context after five generations of selection, (ii) the majority of these DMCs were stably inherited for 2 or 3 generations following the selection experiment, (iii) selection caused overall reductions in epigenetic diversity, and (iv) methylation levels of some CG DMCs were associated with phenotypic changes (Schmid *et al*. 2018).

Genome-wide DNA methylation studies in *Quercus* species showed patterns similar to those obtained in *A. thaliana*: DMCs associated with climate dominate in CG context, and these DMCs occur in or near genes (Platt *et al*. 2015; Gugger *et al*. 2016). Using the experimentally more versatile herb *Plantago lanceolata* as a study organism, Gáspár *et al*. (2019) demonstrated that much of the environment-related epigenetic variation is maintained in an F1 common garden. Thus, at least part of the epigenetic variation observed in the field is stable, non-random and of ecological significance. Although these studies considerably increased our understanding of how epigenetic variation is distributed across large spatial scales, it remains unknown to what extent epigenetic variation contributes to population divergence along small-scale environmental gradients, where the interplay between migration, drift and selection can be extremely dynamic (Richardson *et al*. 2014). Highly heterogeneous environments may thus give rise to distinct signatures of epigenetic variation.

Evidence is accumulating that an epigenetic memory may be particularly beneficial where genetic diversity is in short supply, e.g. following demographic bottlenecks or in clonal plant species (Latzel *et al*. 2016; Ardura *et al*. 2017; Artemov *et al*. 2017; Thorson et al. 2017; Rendina González et al. 2018; Wibowo *et al*. 2018). More fundamentally, Dapp *et al*. (2015), using epigenetic inbred lines of *Arabidopsis thaliana*, demonstrated that epigenetic diversity can drive hybrid vigour in the absence of genetic diversity. Thus, epigenetic variation may be a crucial element of population persistence where evolutionary trajectories or life history traits limit genetic diversity. Study systems combining strong evolutionary pressure (e.g. expansion fronts or heterogeneous environments) and life history traits that constrain genetic diversity (e.g. asexual reproduction and high levels of self-compatibility) are therefore promising for obtaining more insights in the role of epigenetic variation in adaptation and population persistence.

Here, we explore genome-wide epigenetic profiles of F1 plants originating from three natural *Fragaria vesca* populations that were found to harbor strong natural differentiation in terms of traits related to fitness, most likely driven by local topography impacting local soil moisture levels (*De Kort et al*. 2019). Among the most notable patterns observed in a controlled common garden environment were that plants adapted to the stressful conditions at high altitudinal, south-oriented locations were small and produced less flowers than plants originating from low altitudinal, north-oriented locations (*De Kort et al*. 2019). *F. vesca* has also been shown to harbor limited genetic diversity across its range (Hilmarsson *et al*. 2017), presumably as a result of its life history (self-compatible and clonal). Due to this limited genetic diversity in combination with pronounced altitude-dependent phenotypic divergence, we hypothesize substantial adaptive epigenetic signals coinciding with increased stress levels along the studied altitudinal gradient (Fig. 1C). We specifically answer the following questions: (i) do epigenetic memories diverge with altitude, and does methylation increase or decrease with increasing altitude?; (ii) are these small-scale genome-wide methylation patterns comparable to those obtained at a much larger spatial scale (>500 km)?; (iii) are altitudinal DMCs enriched for ecologically relevant gene ontology terms?; and (iv) does acute drought stress induce a detectable epigenetic memory?

**Fig. 1.**
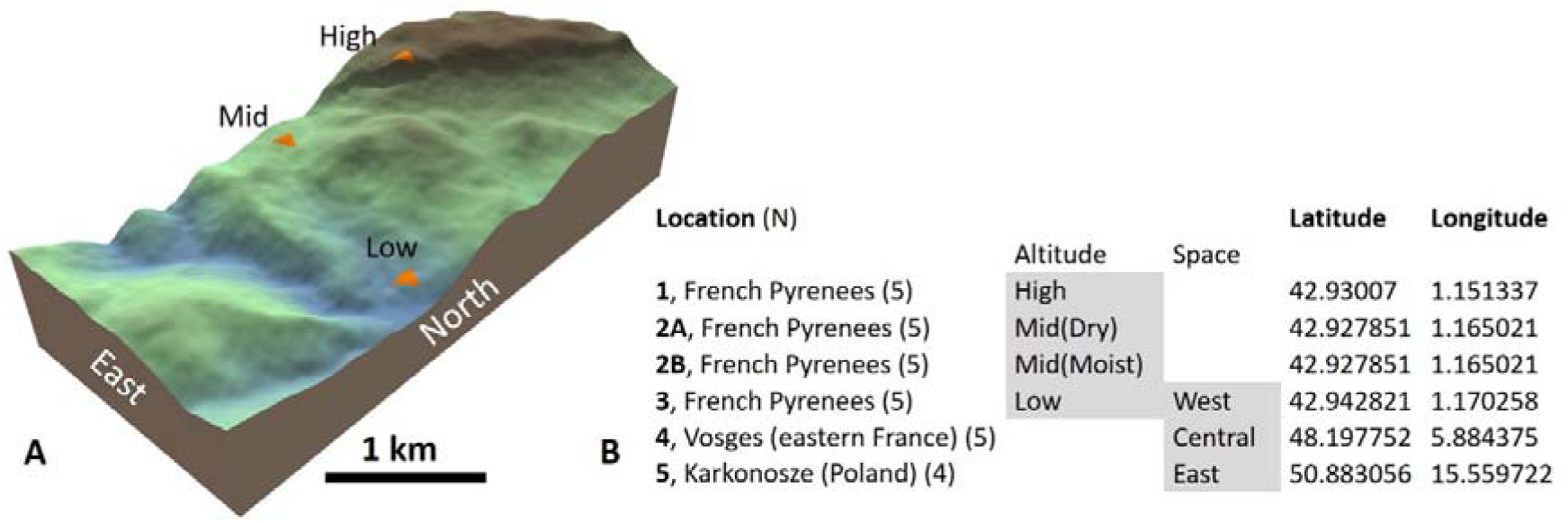
Geographical location of altitudinal WGBS samples (A,B) and of the broader spatial WGBS (B). Samples originate from south western France (Pyrenees), eastern France (Vosges) and south western Poland (Karkonosze). N represents the number of successfully sequenced samples (one Polish sample failed post-sequencing quality checks). Samples from location 2 are used for comparisons within the altitudinal gradient (Mid) as well as between the soil moisture treatments (Dry vs. Moist). Samples from location 3 are used for comparisons within the altitudinal (Low) as well as the spatial (West) gradient. High, mid and low altitude correspond to 1200, 750 and 450 meter asl, resp.

**Fig. 1.**
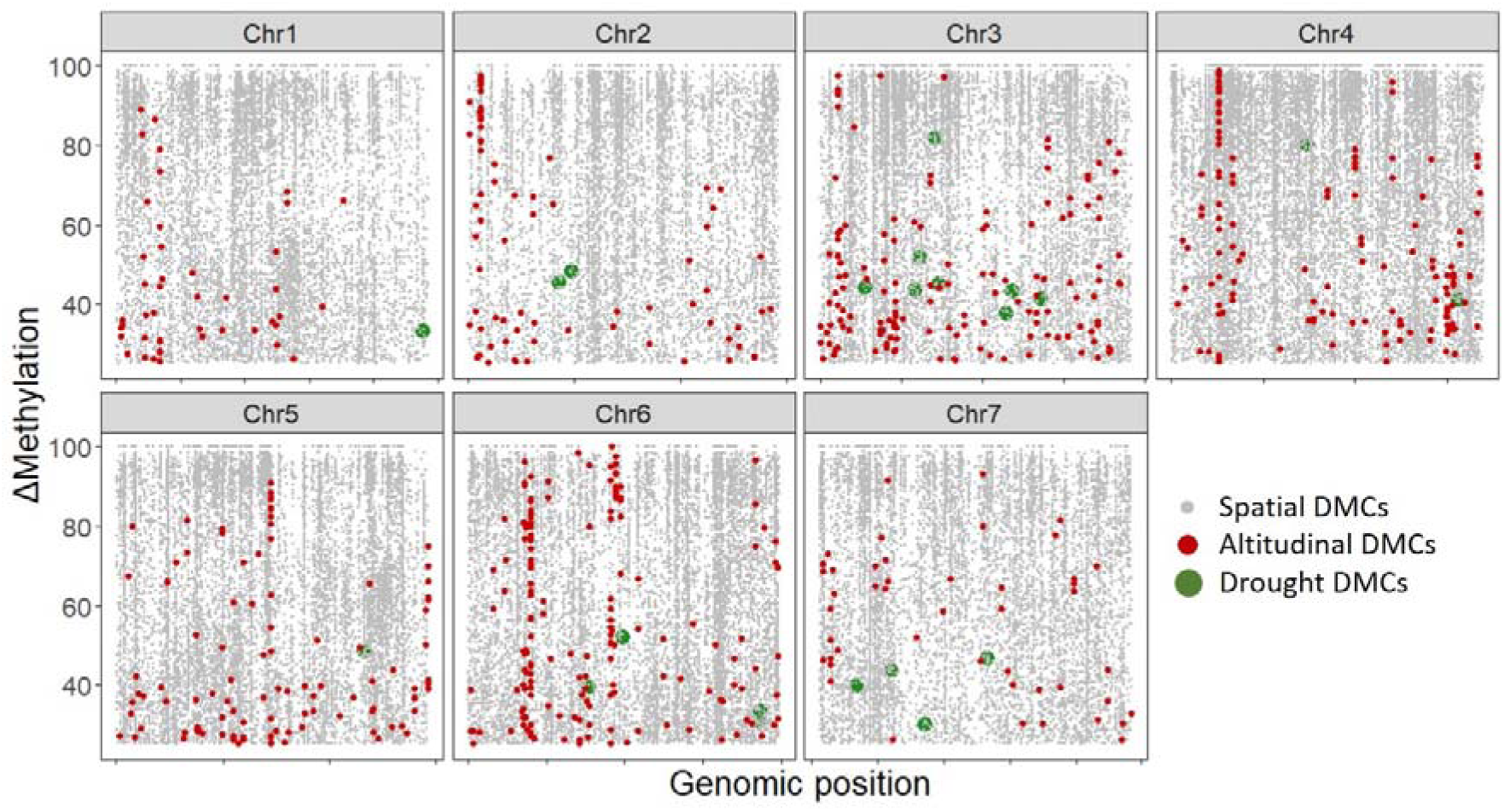
Genome-wide differentiation in methylation for the DMCs detected along the spatial gradient (grey), along the altitudinal gradient (red), and between soil moisture treatments (green).

## METHODS

### Sample collection

Seeds were collected from five plants at (i) three nearby locations in the French Pyrenees, (ii) one location in the French Vosges, and (iii) one location in Poland (Fig. 1). After germination, one seedling per mother plant was randomly selected from every location and grown in humid soil. To compare the magnitude of inherited epigenetic memories to intra-generational epigenetic change acquired through acute drought stress, an additional seedling per mother plant was raised for the mid-altitudinal Pyrenean plants, and these seedlings were subjected to reduced soil moisture levels starting two months after germination. Specifically, watering stopped until leaves went limp, and this process was repeated consecutively for four weeks, after which the plants were allowed to rehydrate during one week to remove most drought-induced epigenetic effects that do not result in a relatively stable epigenetic memory. DNA was then extracted from one leaf per plant, resulting in 30 samples (Fig. 1).

### Whole genome bisulfite sequencing and DMC calling

DNA of 30 freeze-dried samples was extracted with a QIAGEN kit. Up to 200ng of DNA was fragmented to 400bp with a Covaris S2 sonicator prior to whole-genome library preparation (NEBNExt Ultra II kit), ligation of methylated adaptors and size selection on 2% E-gel (450-650 bp). Bisulfite conversion was performed using the EZ DNA methylation gold kit (Zymo Research, Irvine, CA, USA). An enrichment PCR was performed using KAPA Hifi hotstart Uracil+ mastermix in a 12 cycles PCR reaction. Paired-end 75 bp sequencing of the library fragments was performed on eight lanes of an Illumina Hiseq4000 sequencer, generating 76,417,704 ± 13,837,490 reads (mean ± standard deviation) per sample. An extensive quality control was performed on the sequencing data using FastQC version 0.11.5 (Babraham Bioinformatics), showing a mean quality value across each base position in the reads, average quality scores of the reads and average GC content of reads that is to be expected from a high-quality Illumina sequencing run (see Supporting Figs. 1-5 for quality and coverage stat). Trimmomatic (Bolger *et al*. 2014) version 0.36 was used to trim reads for sequences with a Phred score lower than 33 and sequences corresponding to Illumina TruSeq adapters. Sequences shorter than 50bp after trimming were discarded. One sample from Poland (“East”) showed an increased level of duplicate reads and was excluded from further analysis (Supporting Fig. 3).

FastQ Screen version 0.11.1 (Babraham Bioinformatics) was used to check whether the library is consisting of *F. vesca* genomic sequences to rule out contamination with DNA originating from genomes of other species. The trimmed reads were mapped against the Fragaria_vesca_v4.0.a1 genome using bismark version 0.17.0. (Krueger and Andrews 2011). One sample from Poland (“East”) showed suspicious duplication levels and was expelled from further analysis (see Supporting Figs. 1-5 for quality and coverage stats). Average sequencing depth after mapping and deduplication was 30x.

The resulting mapped dataset was used for downstream analyses using R package methylKit version 1.10.0 (Akalin *et al*. 2012). Only CpGs with at least 5x coverage in at least 3 samples per group were retained (Walker *et al*. 2015; Wan *et al*. 2016). To reduce bias due to outlier depth, bases with a read depth above the 99.9th percentile of coverage are filtered out. The filtered data were used to test for differentially methylated CpGs (DMCs), considering a 25% difference and q-values <0.01 as significant. Significant differentially methylated cytosines (DMCs) were identified between (i) low, mid and high altitudinal samples (hereafter “altitudinal DMCs”), (ii) the three distance European samples (hereafter “spatial DMCs”), and (iii) the two soil moisture treatments (hereafter “drought DMCs”).

### Methylation profiling of topography, space and drought

All DMCs were assumed to be ecologically divergent (i.e. resulting from drift or fitness differences). Because the epigenetic signals observed here have persisted in the second generation (F1), they may represent stable epigenetic changes. However, because we cannot corroborate the long-term transgenerational stability of these DMCs, we further refer to ecological rather than evolutionary divergence of methylation patterns. Cytosines that were significant along the altitudinal gradient, were thus considered to be ecologically divergent and potentially adaptive. On the other hand, spatially divergent cytosines that were not significant along the altitudinal gradient, were assumed to be ecologically neutral along the altitudinal gradient. Similarly, the altitudinal DMCs that were not significant along the spatial gradient were assumed to be neutral along the spatial gradient, while spatial DMCs were considered divergent and potentially adaptive along the spatial gradient. The drought DMCs were not significant along the two gradients and were thus considered neutral along both gradients.

As a measure of epigenetic structure (ES), a principal component analysis (PCA) was performed to reveal to what extent DMCs clustered samples according to topography, space and drought. As a measure of epigenetic diversity (ED) that is insensitive to sample size (Vellend *et al*. 2010; Schmid *et al*. 2018), we calculated the average pairwise methylation difference between the five individuals originating from each location. High ED thus reflects high variation in the degree of methylation within a population. Because ED represents a proportion, a quasibinomial model (logit link) was used to test whether ED differs significantly between populations. The resulting effect size (R²) was estimated using the R package “Rsq” (default function), and pairwise comparisons were assessed Tukey-wise (R package “Multcomp”). Methylation shifts, ED and ES were examined for ecologically neutral and potentially adaptive DMCs separately.

### DMC enrichment analysis

Gene ontology terms (GOs) were retrieved from the Genome Database for Rosaceae (GDR, Jung *et al*. 2019). To test which biological processes where overrepresented in sequences containing DMCs (as compared to the full *F. vesca* genome), we performed a Fisher’s exact test with FDR correction as implemented in OmicsBox, using altitudinal DMC GOs as test data and *F. vesca* GOs as reference data. The same analysis was performed for the spatial DMCs, and in CG and non-CG context separately.

To address the role of non-CG methylation in transposon regulation, we aligned genes in which we found one or more DMCs to all transposable elements known in *F. vesca*, using the GDR search function. Specifically, we extracted all genes related to the keyword “transpos” (referring to. e.g. transposase, transposon, transposable element).

## RESULTS

Apart from slight increases in genome-wide methylation levels from high to low soil moisture, and from east to west (Supp. Fig. 6), no systematic genome-wide differences in methylation were observed between the 29 samples. This suggests that the environmental conditions historically encountered by our *F. vesca* populations target specific genomic regions or cytosine sites. We accordingly detected a total of 82 839, 699 and 23 DMCs along the spatial gradient, the altitudinal gradient and between the soil moisture treatments, respectively (Supp. Table 1). These DMCs often clustered together in genomic islands of differential methylation (Fig. 1, Fig. 2A). A Mann-Whitney U-test confirmed this clustering, showing that DMCs in all sequence contexts were significantly more proximate to one another than expected based on the genome size (240 Mbp) and on the genome-wide number of DMCs (varying from 13 altitudinal DMCs in CG context to 51135 spatial DMCs in CG context) (Supp. Fig. 7, Supp. Table 2).

**Fig. 2.**
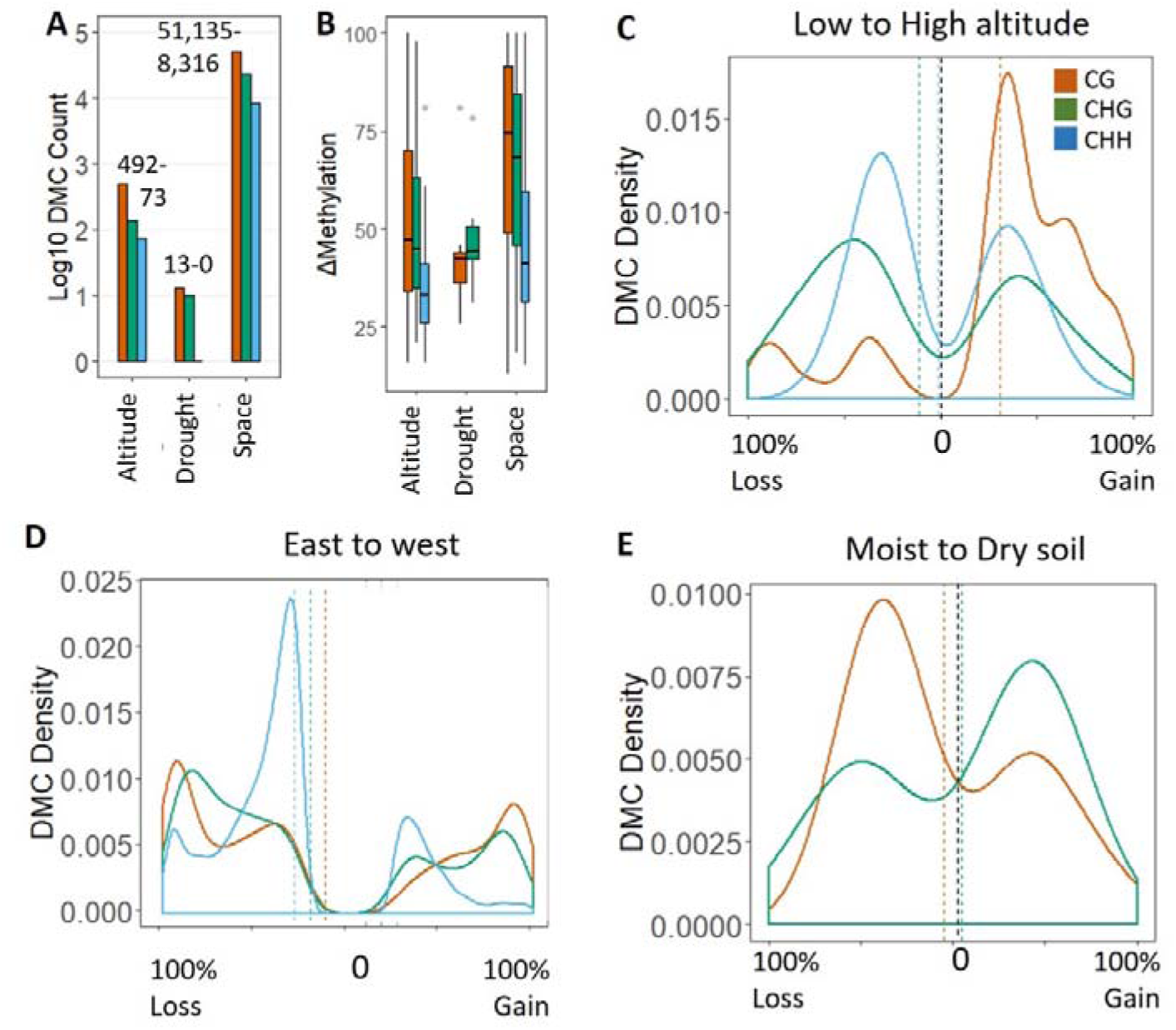
Distribution of methylation patterns among the studied gradients, including DMC counts (A), DMC differentiation (B), and DMC density of change in methylation level along the topographical gradient (C), the spatial gradient (D) and between soil moisture treatments (E). All patterns were visualized for each sequence context separately (CG in orange, CHG in green en CHH in blue).

Methylation divergence was most pronounced along the spatial gradient, followed by the altitudinal gradient and the soil moisture treatments (Fig. 2B). A total of 59 altitudinal DMCs (8.5%) were also significantly divergent along the spatial gradient, and another 153 altitudinal DMCs (21.9%) were located within 500 bp of a spatial DMC, indicating shared methylation patterns among distinct spatial scales. The negligible proportion of genome-wide cytosines that was differentially methylated between soil moisture treatments suggests that short-term acute soil dryness does not constitute a pronounced epigenetic memory.

DMC density profiles were substantially different between the altitudinal gradient, the spatial gradient and the soil moisture treatments (Fig. 2C-E). For altitudinal DMCs, the most dominant shift in methylation was observed in CG context, with substantial methylation gain as altitude increased (Fig. 2C). However, the most extreme shifts in DMC methylation level (i.e. toward 100% methylation loss or gain) was observed along the spatial gradient (Fig. 2D). Thus, fixation of methylation patterns occurred both at small and large spatial scale, but was more frequent along the spatial gradient.

A total of 247 out of 698 DMCs (35.4%) systematically gained (113 CG, 21 CHG and 20 CHH) or lost (38 CG, 35 CHG and 20 CHH) methylation from low to high altitude. Methylation gains along the altitudinal gradient thus predominantly occurred in the CG context (see also Fig. 2C). Along the spatial gradient, a total of 56 795 out of 82 839 DMCs (68.6%) systematically lost (18 099 CG, 8731 CHG and 3609 CHH) or gained (17 038 CG, 7680 CHG and 1638 CHH) methylation from east to west (see also Fig. 2D). The plants from western Europe thus particularly differ from central Europe in the amount of demethylating CHH sites. Given that the plants raised here were genetically variable, we assume that the observed drastic and systematic shifts in methylation from low to high altitude and from east to west are (i) not solely driven by genetic differences, and/or (ii) underpinned by epistatic effects.

Methylation patterns grouped the plants according to their location of origin (Fig. 3, Supp. Table 3), thus convincingly indicating that diverging methylation levels persisted through the F1 generation. Our results further show that neutrally behaving cytosines can align with spatial organization, particularly at large spatial scales (Fig. 3B, 3D), as expected under isolation-by-distance. This finding is concordant with the behavior of neutral genetic markers that spatially cluster as a result of drift or gene flow. Clustering was more pronounced for adaptive than for neutral DMCs (Fig. 3A, 3C), indicating epigenetic isolation-by-ecology.

**Fig. 3.**
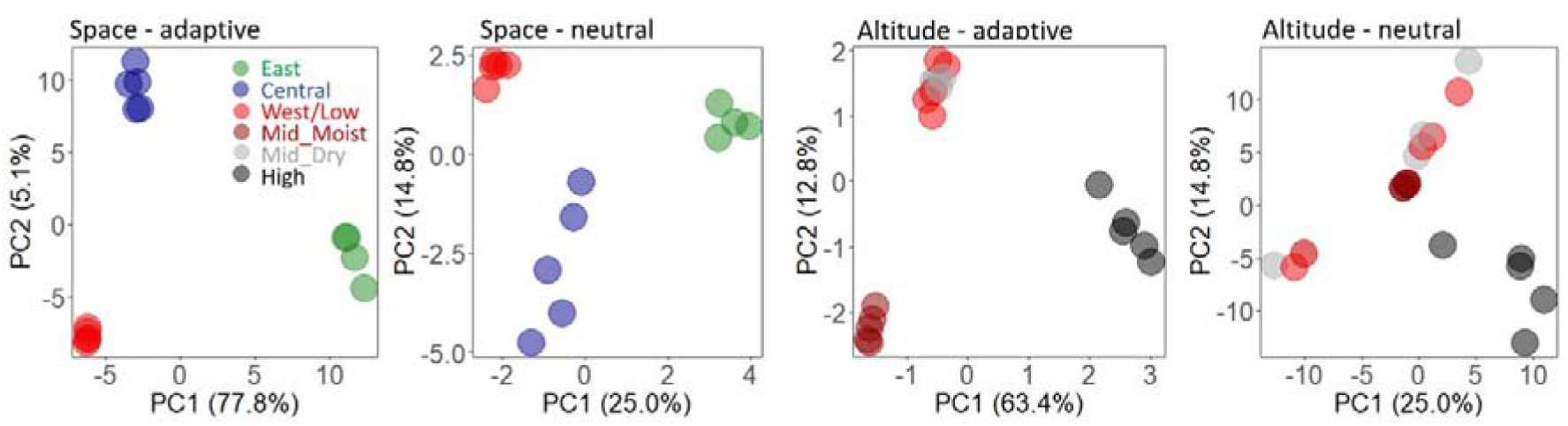
Principal components analysis of methylation levels at large and small spatial scale, and for ecologically neutral vs. potentially adaptive methylation marks. See supporting Table 3 for eigenvalues and variable contributions.

Adaptive epigenetic diversity (ED) significantly increased from low to high altitude in CG context, but not in non-CG context (Fig. 4A-B, Supp. Table 4). There also was a significantly lower adaptive ED in the west than in the center and east of the sampling area, both in CG and non-CG context (Fig. 4A-B, Supp. Table 4). As opposed to adaptive ED, neutral ED did not significantly change with altitude or space (Fig. 4C-D, Supp. Table 4).

**Fig. 4.**
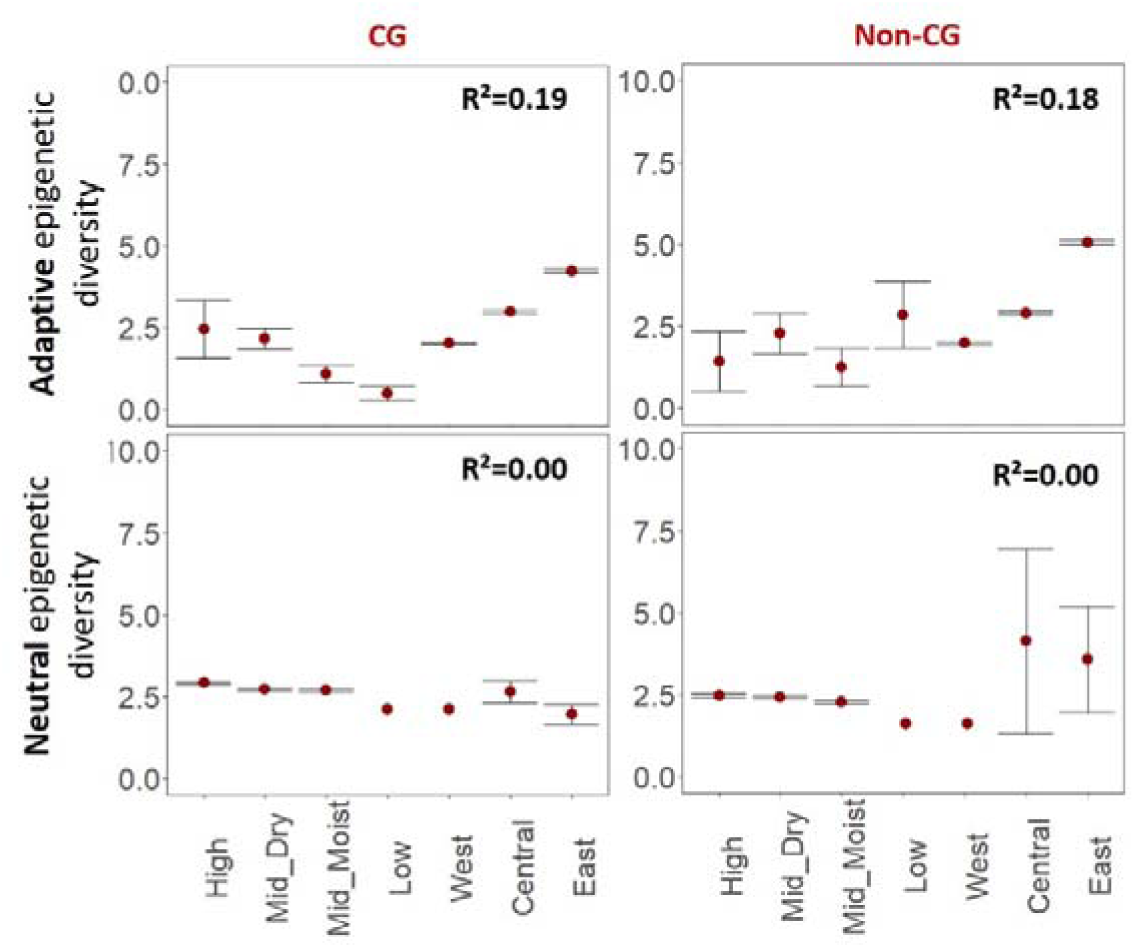
Epigenetic diversity (ED) for ecologically neutral vs. putatively adaptive methylation, both in CG and non-CG context. Effect sizes are shown as R² retrieved from generalized linear models testing for differences in ED between locations. Error bars represent 95% confidence intervals of the means. See Supp. Table 4 for corresponding statistics.

Out of 138 and 25 GO terms that were associated with the altitudinal DMCs in CG and non-CG context, respectively, 54 (39.1%) and 2 (8.0%) were significantly enriched in comparison to the full *F. vesca* GO compilation (Supp. Table 5, 6). A similar proportion of enriched GO terms was observed for the spatial DMCs in CG context (39.7%, i.e. 578 out of 1456 GOs). As opposed to the altitudinal DMCs, however, a high proportion of enriched GOs was also found in non-CG context (48.0%, i.e. 290 out of 604 GOs).

DMCs in CG vs. non-CG context represented a distinct set of enriched biological processes (Fig. 5). Where non-CG DMCs were particularly enriched for regulatory functions (e.g. regulation of gene expression and protein dephosphorylation), biological processes more directly related to environmental stressors were overrepresented only in CG-context (e.g. circadian rhythm, antibiotic metabolism and response to light). GOs related to cell division and reproduction (e.g. spindle organization and sexual reproduction) were enriched in both CG and non-CG DMCs. No pronounced differences in GO composition were found between altitudinal and spatial DMCs, and most altitudinal GOs (69%) were also part of the spatial GO distribution (Supp. Table 5).

**Fig. 5.**
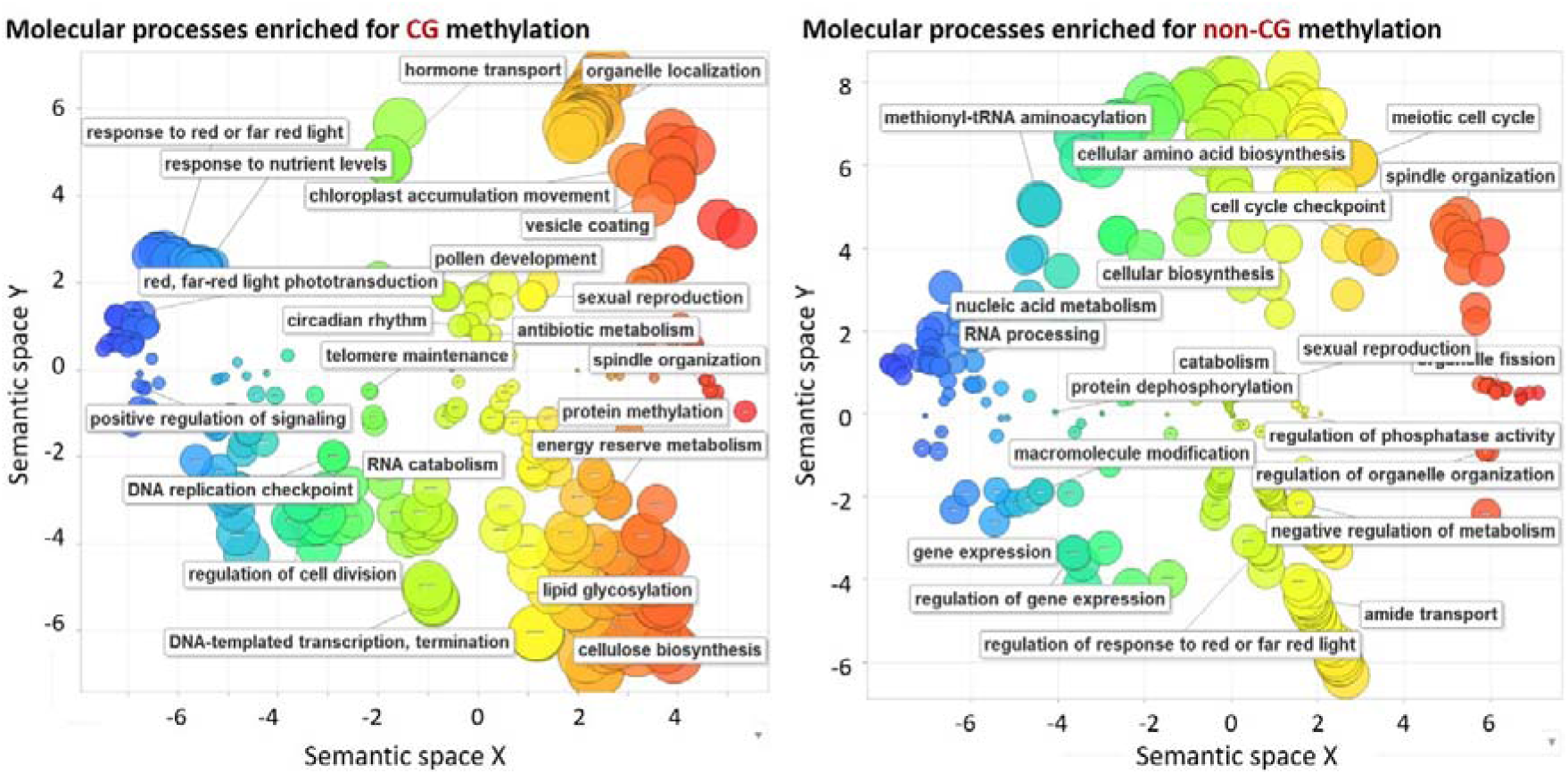
GO enrichment graphs showing the significantly enriched GO terms in CG (Left) and non-CG context for the spatial DMCs. Terms were visualized using REVIGO (Supek *et al*. 2011), which cluster GO terms according to their semantic similarities (e.g. response to abiotic stimulus and response to stimulus are clustered together). In each cluster, one or two GOs representing their corresponding cluster are shown. These graphs represent enrichment in spatial DMCs. Please see Supp. Table xx for GOs enriched in altitudinal DMCs.

A total of 31 transposable elements were found to contain one or more DMCs (Supp. Table 1). The most heavily differentiated transposon harbored no less than 21 DMCs, of which 15 in non-CG and 6 in CG context. All but one differentially methylated transposon were found at the large spatial scale, and were dominated by DMCs in non-CG context (Fig. 6). On average, transposon DMCs lost 18% and 32% of methylation from east to west in CG and non-CG context, respectively.

**Fig. 6.**
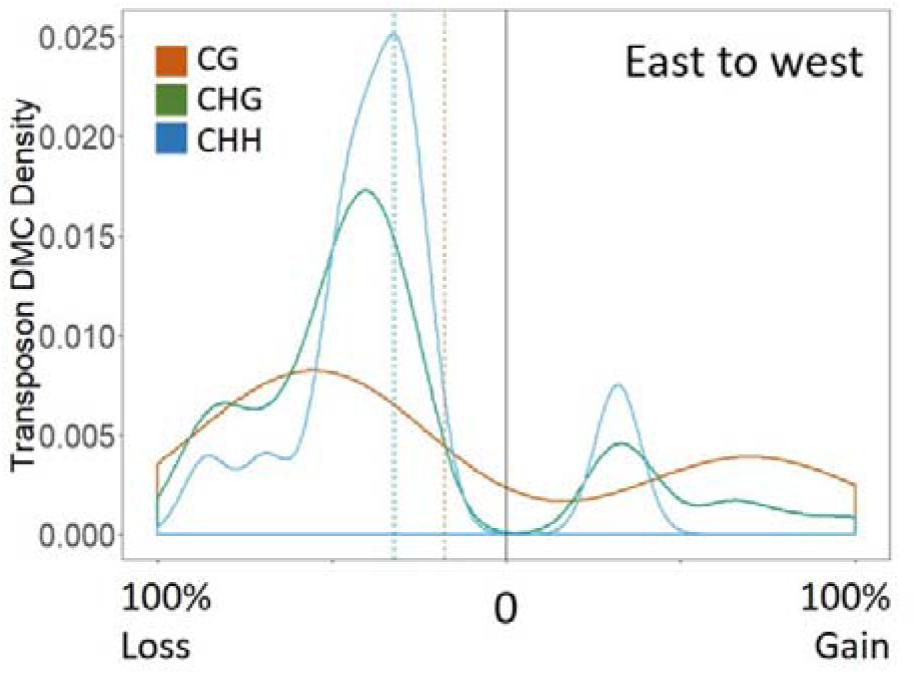
DMC density plot showing methylation gain and loss for transposable elements, in CG, CHG and CHH context.

## DISCUSSION

Epigenetic variation in natural populations probably is key to their survival, particularly when they are genetically depleted and environmentally challenged. In such systems, it may be favorable for individuals to acquire an epigenetic memory that allows efficient responses to fluctuating environmental stressors. Here, we shed light on the prevalence of natural epigenetic variation along a steep environmental gradient, and put these findings into a much wider geographical context. Collectively, our results indicate that epigenetic memories develop both at small and large spatial scales, each associated with distinct epigenetic signatures. Specifically, methylation in non-CG context gains in importance as the spatial scale increases, and this translates into more methylation differentiation of regulatory sequences and transposable elements. Conversely, divergence of CG methylation was more pronounced at the fine-scale altitudinal gradient, where it may guide adaptive gene expression in response to environmental variability.

We found that epigenetic memories, predominantly in CG context, can diverge even at very small spatial scale (< 2 km), indicative of epigenetic isolation-by-ecology. This finding is in strong contrast to an earlier study that was unable to detect significant epigenetic differences between alpine herb populations originating from three elevations and grown in a common garden using 150 methylation-sensitive AFLPs (MS-AFLPs) (Nicotra *et al*. 2015). In our study, methylation in CG-context increased from low to high altitude, suggesting that high altitudinal environments trigger activation of genes predominantly underpinning amino acid metabolism, intracellular transport, responses to light conditions and cellulose catabolism (Supp. Table 6). Furthermore, we observed a pronounced increase in epigenetic diversity for CG DMCs from low to high altitude, suggesting that epigenetic diversity for ecologically adaptive methylation sites may be favorable where the environment is more heterogeneous. While evidence supporting DNA methylation differentiation in response to environmental heterogeneity is still lacking, several lines of evidence have demonstrated increased up-regulation of genes involved in environmental responses to novel and/or stressful conditions through gene body methylation (Artemov *et al*. 2017; Dixon *et al*. 2018).

Fine-scale epigenetic patterns differed substantially from large-scale epigenetic patterns. As compared to the fine-scale altitudinal gradient, the large spatial gradient was featured by (i) an increase in the number of DMCs with two orders of magnitude, (ii) more intense methylation differentiation (on average 70% vs. 40%), (iii) a more prominent role for non-CG differentiation and differential transposon suppression, and (iv) more pronounced population structure, particularly for ecologically adaptive DMCs. Although the CG-context constituted the most divergent methylation patterns irrespective of spatial scale (Fig. 2), the proportion of non-CG DMCs increased considerably from small to large spatial scale (Fig. 2D). This finding is in agreement with a large-scale study on *A. thaliana* showing that non-CG demethylation associated with transposon activity was abundant where temperature reached extreme levels (Keller *et al*. 2016). It is unclear, however, whether transposon activity was more related to temperature extremes than to demographic history and range dynamics, given that highest transposon activity was observed at *A. thaliana*’s range edges where its distribution becomes more scattered (Beck *et al*. 2008; Alonso-Blanco *et al*. 2016). Here, non-CG DMCs lost methylation and were less diverse (Fig. 4, 6) from east to west, which, hypothetically, results from ecology-driven activation of transposons towards the edge of the distribution range of *F. vesca* (defined by the Pyrenees in the southwest of its distribution), where transposon activation through demethylation may provide opportunities for genetically impoverished populations to boost genetic change. Although this would be in line with earlier studies pointing to an evolutionary rescue mechanism for transposons during range expansions (Stapley *et al*. 2015; Rey *et al*. 2016b), more research on the spatial distribution of transposon activity and its role in evolution is required to validate this assumption. Nevertheless, the observation that transposons are differentially suppressed at large spatial scale and not along a fine-scale steep gradient, suggests that differential suppression of transposon activity follows biogeographic storylines rather than fine-scale environmental clines.

The more pronounced epigenetic clustering for neutrally behaving cytosines along the spatial gradient than along the altitudinal gradient (Fig. 3) may suggest epigenetic isolation-by-distance (Whipple & Holeski 2016). However, most clustering occurred at putatively adaptive DMCs indicating a more dominant role for isolation-by-ecology over isolation-by-distance. Given that roughly 30% of altitudinal DMCs were shared with the spatial DMCs, part of the adaptive epigenetic divergence along both spatial scales may be driven by parallel ecological processes. This spatial ecological parallelism underlying methylation patterns is corroborated by the strong overlap in enriched gene ontology processes between both spatial scales (Fig. 5, Supp. Table 5, 6).

While the origin-dependent epigenetic memories (i.e. altitudinal and spatial DMCs) were stably transmitted to the F1 generation, acute drought stress-induced epigenetic signatures were weak (Fig. 1,2). This suggests that repeated exposure to stressful conditions is required for acquiring a detectable epigenetic memory, and emphasizes the importance of historical stress experience for the generation of an epigenetic memory. Vice versa, our results suggest that the loss of an epigenetic memory requires long-term release of stressful conditions, and that multiple generations without stress exposure are required for completely resetting the epigenetic machinery. Multi-generational persistence of epigenetic signatures (i.e. epigenetic carryover) and thus slow trans-generational loss of epigenetic memory is a typical epigenetic mechanism observed in common gardens quantifying epigenetic variation across generations (Paszkowski & Grossniklaus 2011; Miska & Ferguson-Smith 2016; Proulx *et al*. 2019). Given the natural ubiquity of transgenerational epigenetic inheritance, at least part of the epigenetic patterns observed in our F1 generation is expected to reflect such epigenetic carryover. Nevertheless, the precise extent of multi-generational methylation inheritance requires additional generations of epigenetic profiling.

Collectively, our findings provide novel insights into the natural prevalence of adaptive epigenetic divergence and the processes driving epigenetic memories at distinct spatial scales. We showed that significantly different epigenetic memories, presumably concentrated in or near gene bodies, arise at fine spatial scales. At large spatial scale, epigenetic memories also diverge at the level of regulatory sequences and transposons. We hypothesize that genetic and epigenetic responses complementary support fitness in heterogeneous environments, and that non-CG demethylation increases in importance as genetic variation gets depleted. Further research involving higher resolution sampling and a multi-generational common garden is required to shed more light on the role of epigenetic variation at distinct spatial scales. This would particularly increase our understanding of epigenetic memory acquisition and divergence as an adaptive strategy of natural populations that could enhance their ability to cope with global change stressors.

## ACKNOWLEDGEMENTS

HDK holds a postdoctoral fellowship funded by FWO (Research Foundation Flanders, 12P6517N). Dr. Kenny Helsen and Kasper Van Acker helped raising the plants. Dr. Martin Diekmann and Josef Müller took care of the sampling in Poland.

## AUTHOR CONTRIBUTIONS

HDK coordinated the research, performed data-analyses of DMCs and wrote the manuscript. FVN performed all bio-informatics analyses. MD assisted with field sampling. All co-authors provided comments and suggestions to the first version of the manuscript.

