## Supporting Fig for "Ecological divergence of DNA methylation patterns at distinct spatial scales"

**Supporting Figures 1 - 7**

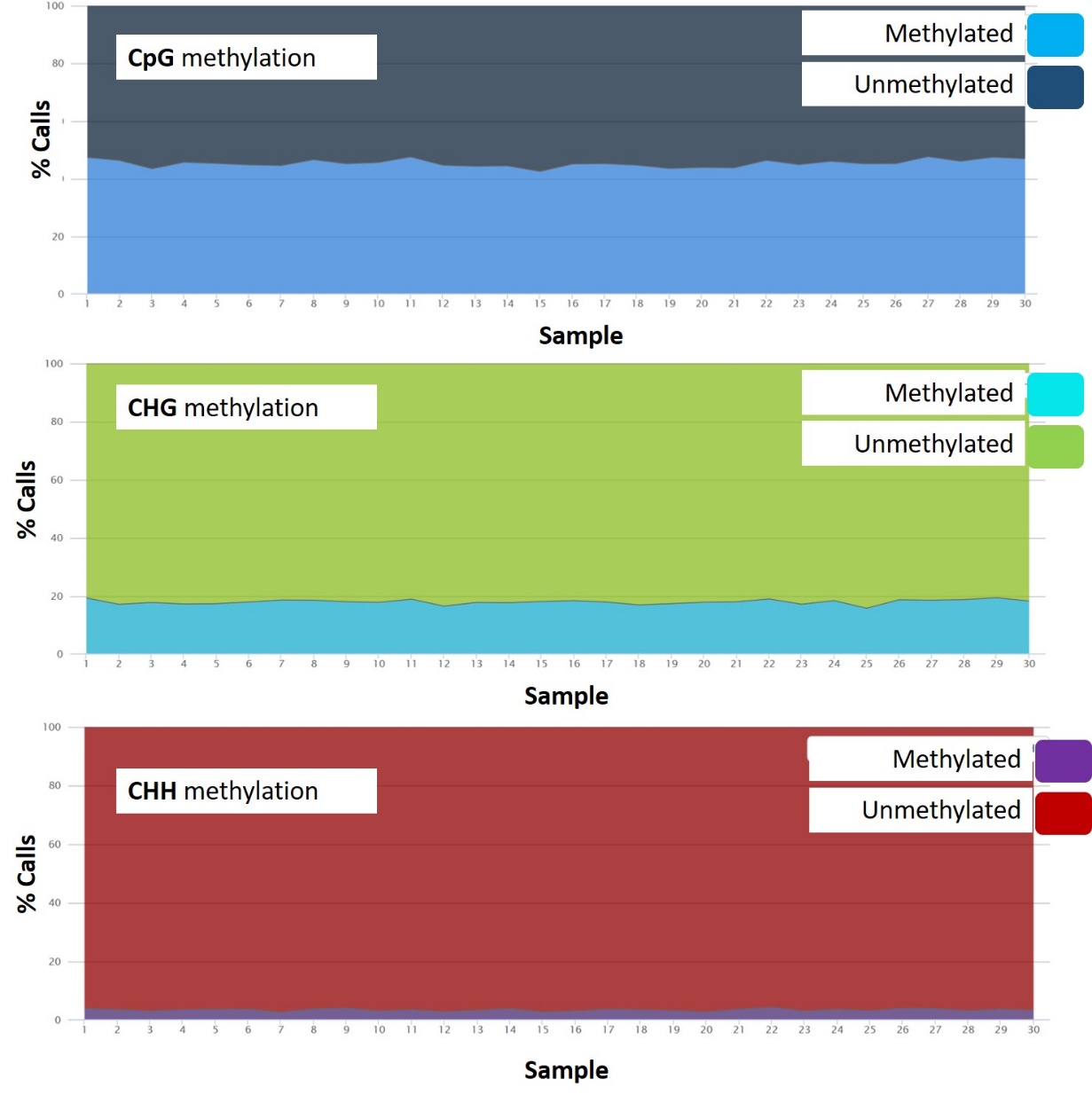

**Supp. Fig. 1**. Proportion of cytosines that were methylated across the genome, within each sample, and for each sequence context (CG, CHG and CHH, respectively).

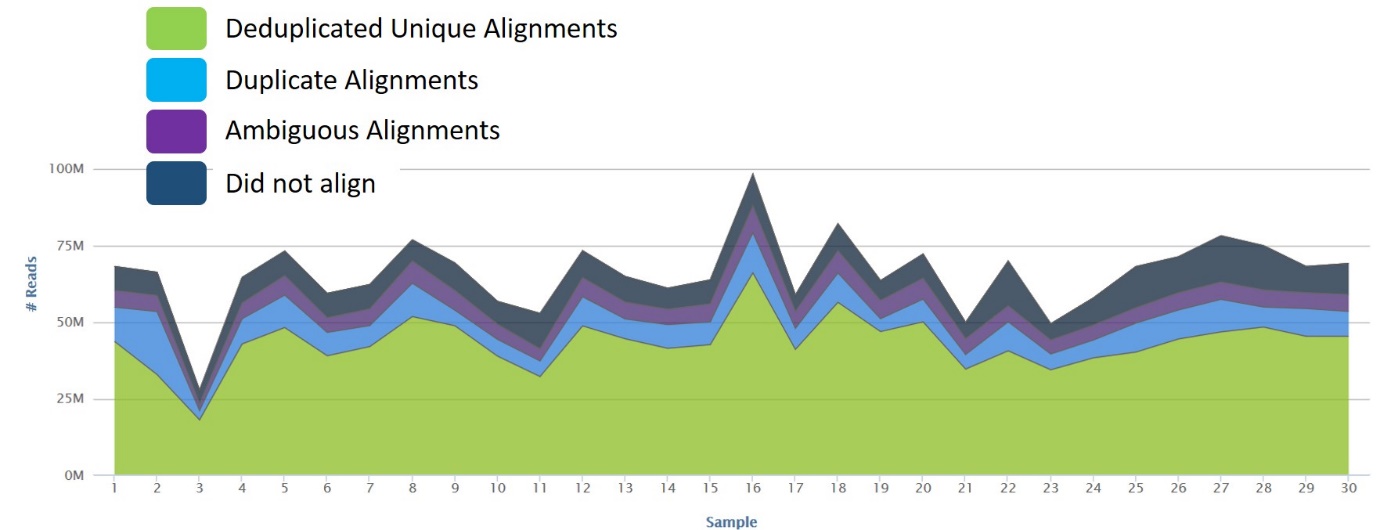

Supp. **Fig. 2.** Number of reads sequenced and aligned against the *F. vesca* genome in each sample.

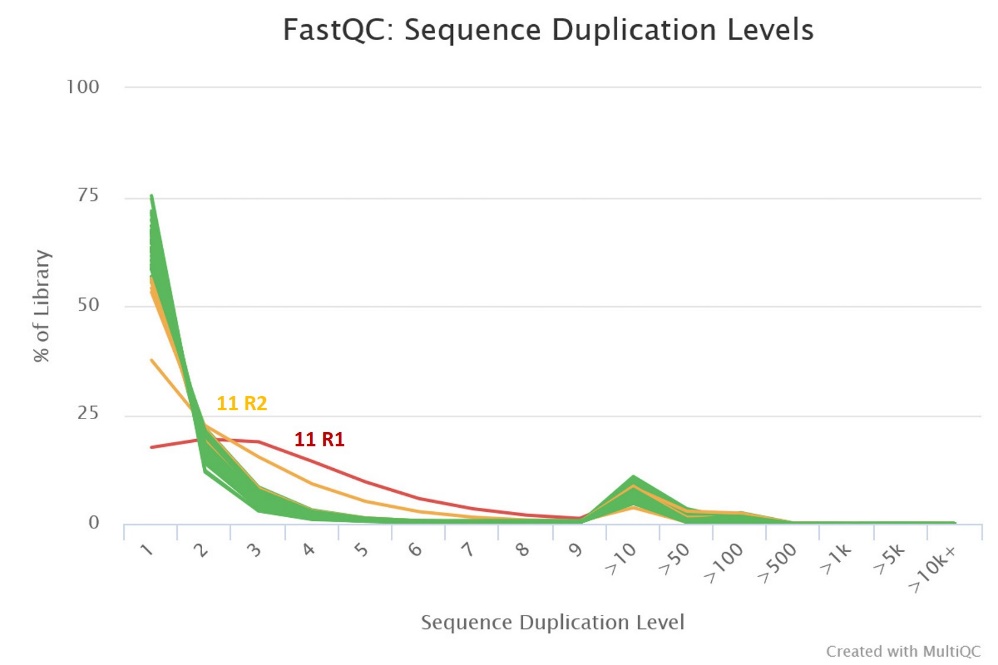

**Supp. Fig. 3**. Sequence duplication levels for each of the 30 individuals subjected to whole genome bisulfite sequencing. Individual 11 showed suspicious levels of duplication and was therefore removed from the data-analysis.

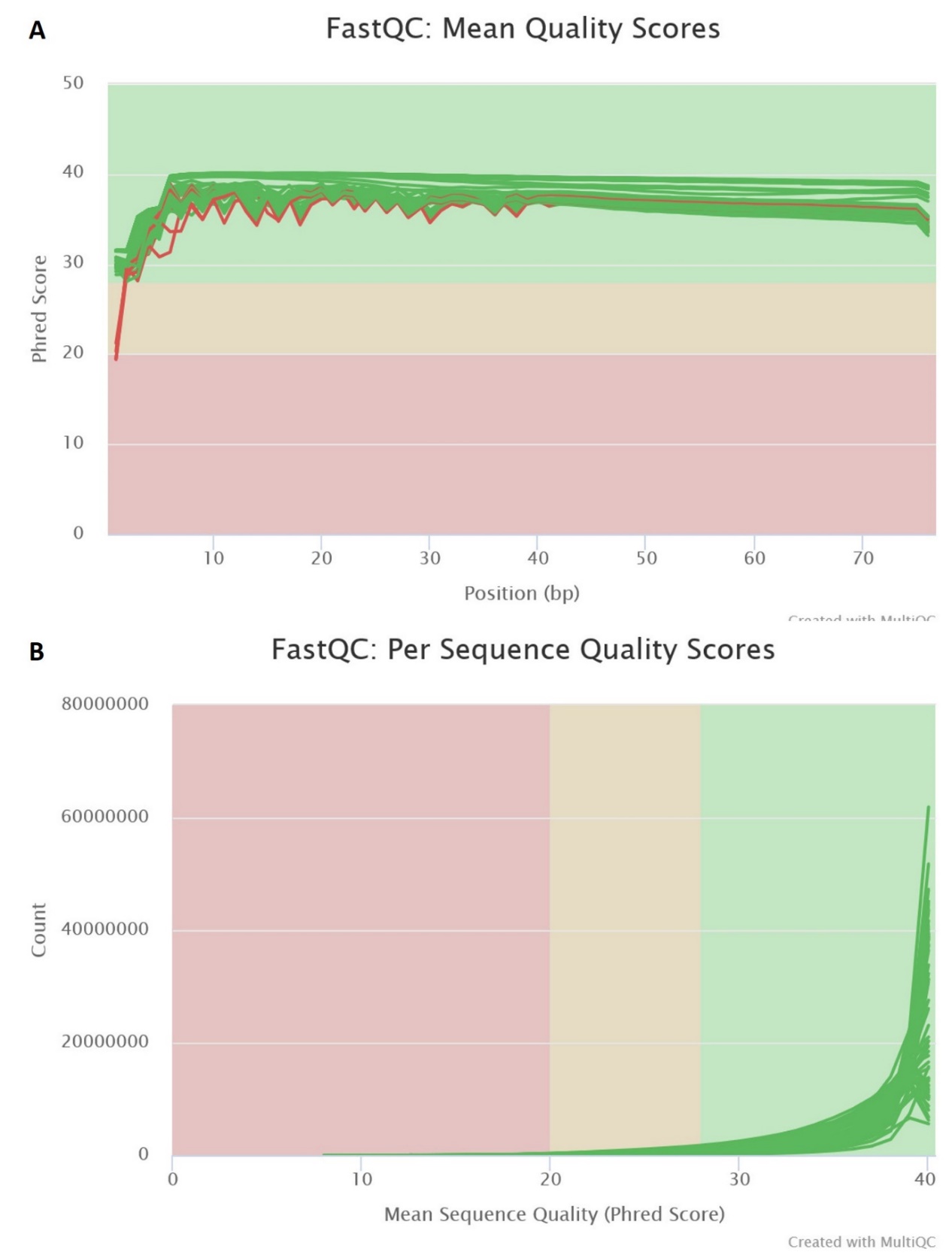

**Supp. Fig. 4**. Per base-pair (A) and mean sequence (B) quality scores of final reads (trimmed to 75 bp) ranged from acceptable to excellent.

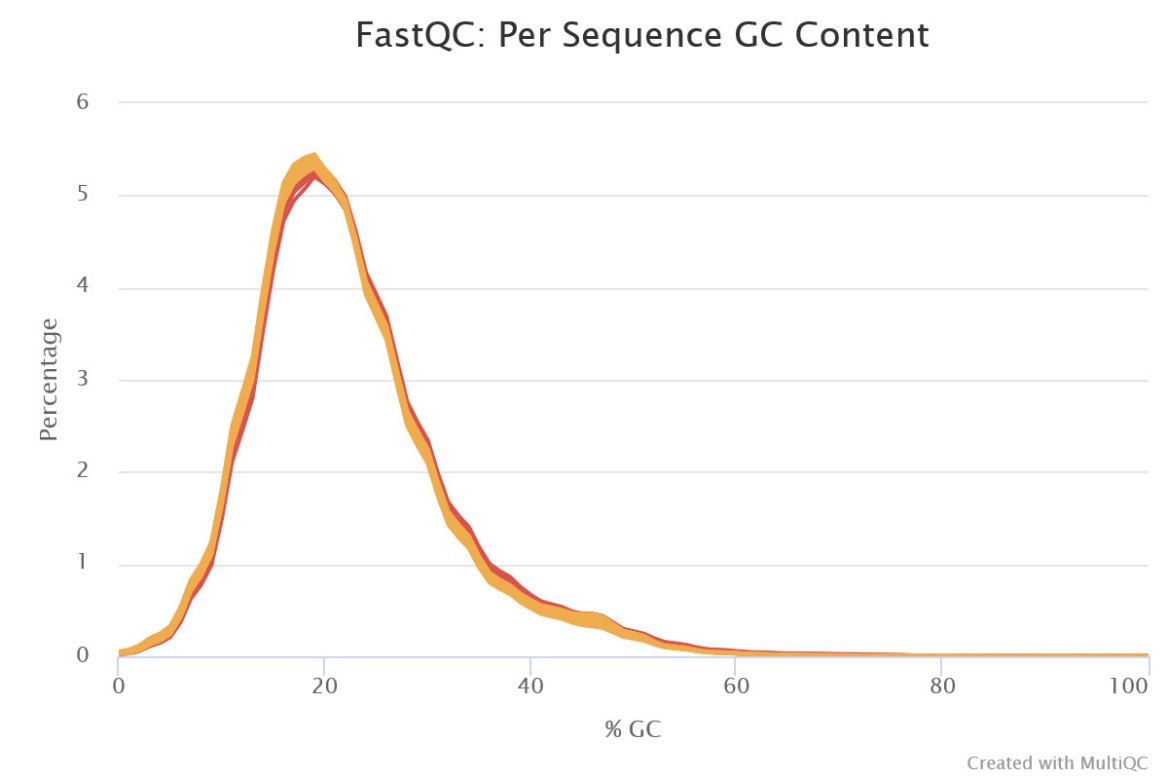

**Supp. Fig. 5.** Per sequence GC content.

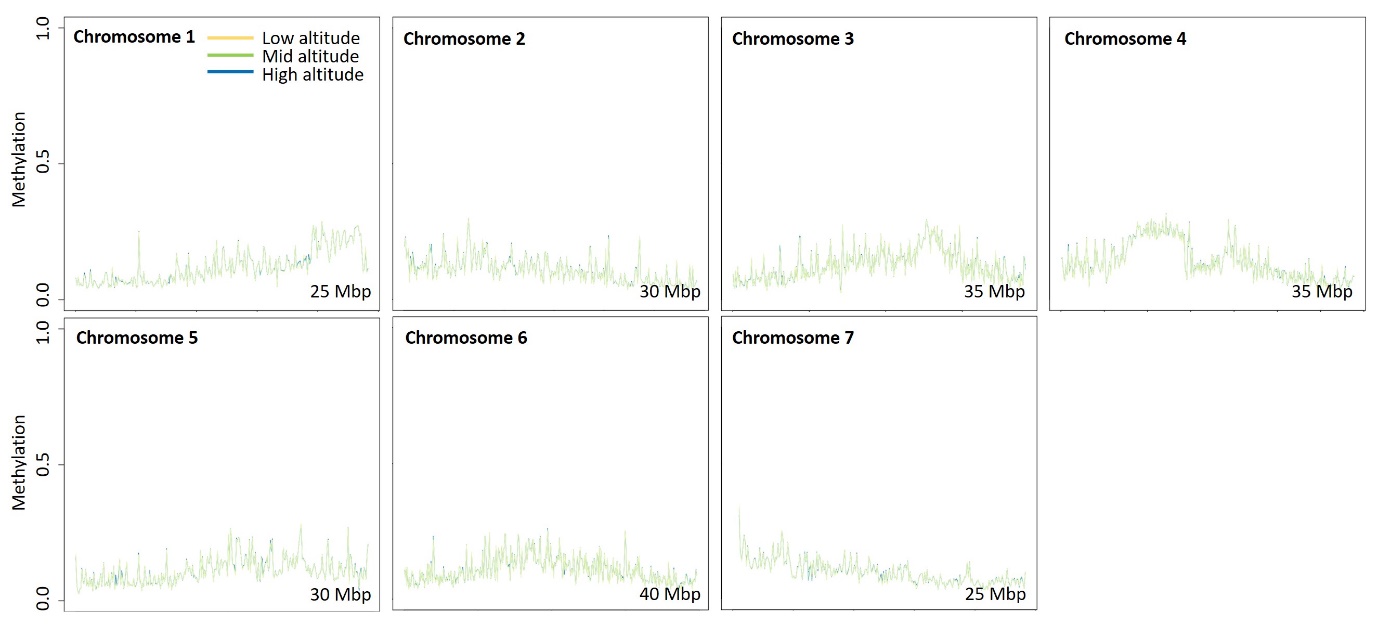

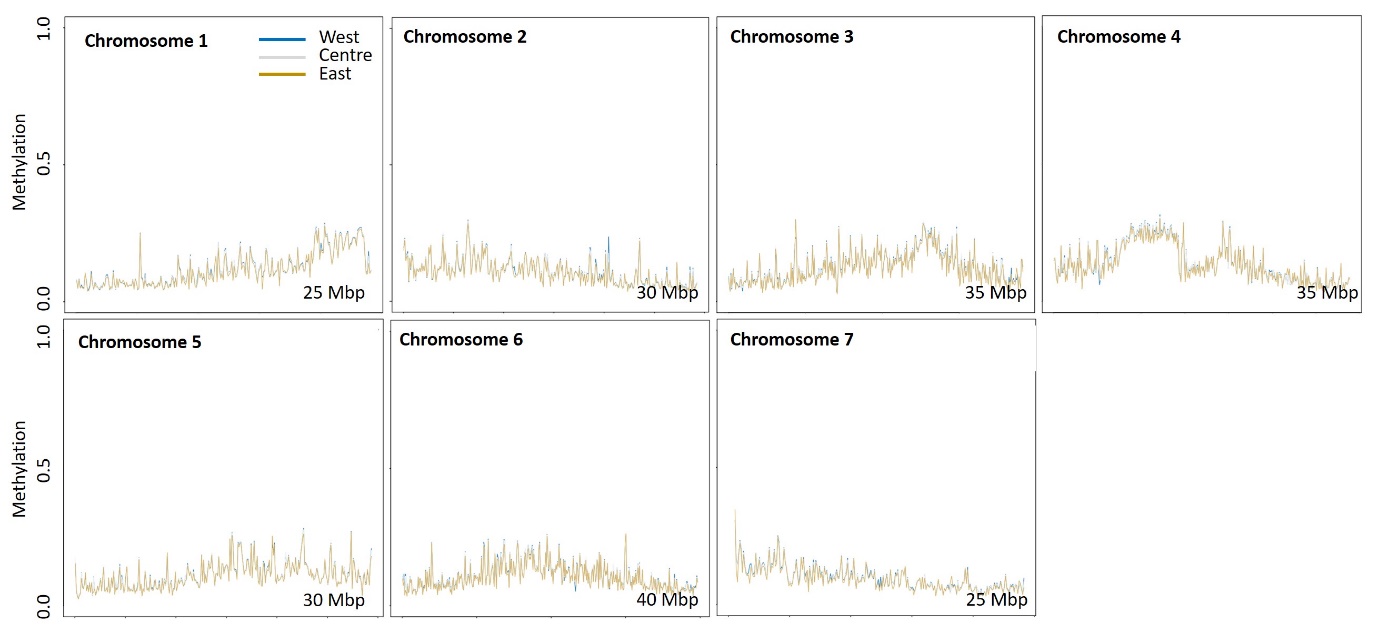

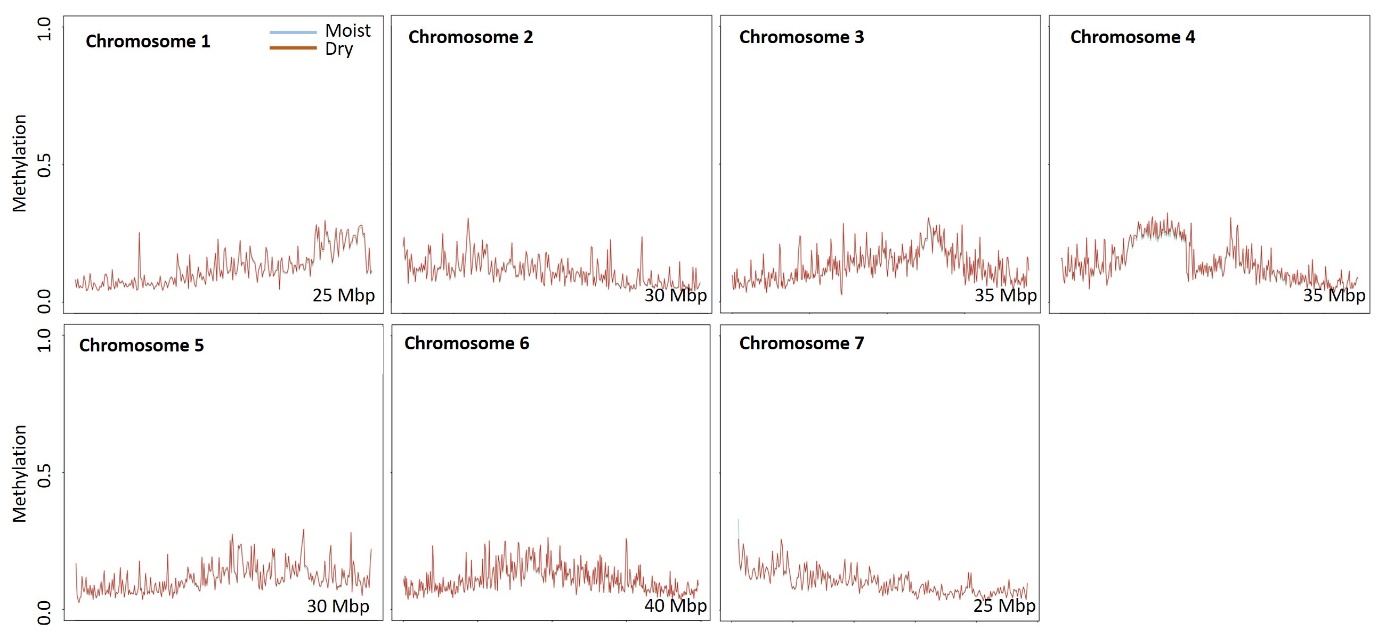

**Supp. Fig. 6.** Genome-wide methylation levels at small spatial scale (upper graphs), large spatial scale (central graphs), and between soil moisture treatments (lower graphs).

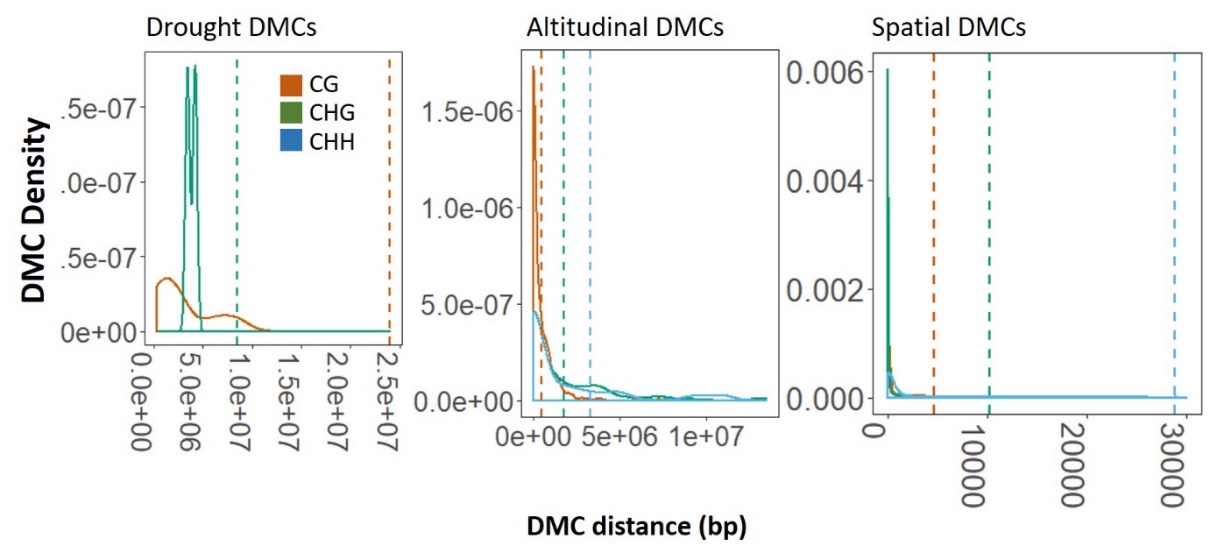

**Supp. Fig. 7.** Density plots showing distribution of pairwise distances between consecutive DMCs. Dashed lines correspond to average expected pairwise distance, expected based on genome size (240 Mbp) and number of DMCs in each context and gradient (see Supporting Table 1 for more details).
